# Metformin treatment results in distinctive skeletal muscle mitochondrial remodeling in rats with different intrinsic aerobic capacities

**DOI:** 10.1101/2024.03.01.582957

**Authors:** Matthew P. Bubak, Arik Davidyan, Colleen L. O’Reilly, Samim A. Mondal, Jordan Keast, Stephen M. Doidge, Agnieszka K. Borowik, Michael E. Taylor, Evelina Volovičeva, Michael T. Kinter, Steven L. Britton, Lauren G. Koch, Michael B. Stout, Tommy L. Lewis, Benjamin F. Miller

## Abstract

The rationale for the use of metformin as a treatment to slow aging was largely based on data collected from metabolically unhealthy individuals. For healthspan extension metformin will also be used in periods of good health. To understand potential context specificity of metformin treatment on skeletal muscle, we used a rat model (HCR/LCR) with a divide in intrinsic aerobic capacity. Outcomes of metformin treatment differed based on baseline intrinsic mitochondrial function, oxidative capacity of the muscle (gastroc vs soleus), and the mitochondrial population (IMF vs SS). Metformin caused lower ADP-stimulated respiration in LCRs, with less of a change in HCRs. However, a washout of metformin resulted in an unexpected doubling of respiratory capacity in HCRs. These improvements in respiratory capacity were accompanied by mitochondrial remodeling that included increases in protein synthesis and changes in morphology. Our findings raise questions about whether the positive findings of metformin treatment are broadly applicable.

---

Metformin is the most prescribed drug for type 2 diabetes (T2D) and is the fourth most prescribed pharmaceutical in the United States. Metformin has over 50 years of safety data, is cost effective, and is associated with a decreased risk of mortality and morbidity for diseases such as cardiovascular disease, cancer, and dementia^1–3^. The beneficial effects of metformin on multiple chronic diseases make it an attractive candidate to slow aging and extend healthspan, which led to the proposed Targeting Ageing with Metformin (TAME) trial^4^. A meta-analysis of clinical trials suggests that people with T2D taking metformin have lower all-cause mortality and cancer incidence than other diabetics and the general population^5^. Much has also been made about the observation that metformin-treated patients with T2D had better survival rates than their matched non-diabetic control group^6^. However, a recent study that followed individuals who take metformin for twenty years showed that T2D patients had shorter survival than controls^7^. The effects of metformin in healthy individuals are less understood.

One accepted definition of healthspan is the period free of chronic disease^8^. Therefore, treatments, like metformin, which prolong healthspan would start while an individual is healthy and absent of chronic disease. We highlighted that it appears that benefits of metformin are reduced in those without chronic disease^9^. In the landmark Diabetes Prevention Program (DPP), secondary analyses showed that metformin had minimal effects in individuals with a lower BMI (<30 kg/m^2^) and fasting glucose (<110 mg/dL)^10^. Furthermore, a small clinical trial showed that metformin increased insulin sensitivity in subjects with T2D or had a family history of T2D, but worsened insulin sensitivity in other subjects^11^. Finally, our previous study^12^ and the MASTERS trial^13^ showed that metformin inhibited the positive effects of aerobic or resistance training, respectively, in older healthy individuals. The absence of a positive effect of metformin treatment in healthy individuals is relatively inconsequential, but potential detrimental effects would render metformin treatment untenable. Thus, the efficacy of metformin to extend healthspan in healthy populations warrants further investigation.

Surprisingly, the mechanisms of metformin are only partially understood. One proposed action of metformin is a disruption of mitochondrial respiratory capacity through inhibition of complex I (CI) of the electron transport system (ETS)^14,15^ with subsequent activation of AMP-activated protein kinase (AMPK)^16^. A second potential mechanism results from the positive charge of metformin, which accumulates in the inner mitochondrial membrane space and disrupts mitochondrial membrane potential^17^. This potential mechanism would result in an inefficiency in energy transfer and active AMPK^16^. Chronic AMPK activation results in mitochondrial remodeling through protein turnover and fission/fusion events^18^ to reestablish energetic homeostasis. Mechanisms of mitochondrial remodeling are energetically costly and result in additional cellular energetic stress. Thus, the ultimate success or failure of the mitochondrial remodeling depends on the energy production and the cellular priorities for energy.

Aerobic capacity (i.e. VO_2max_) is a major predictor of all-cause morbidity and mortality^19,20^. While determinants of aerobic capacity are multifactorial, one important determinant is energy transfer in mitochondria^21,22^. High-capacity and low-capacity running rats (HCRs and LCRs, respectively) are outbred lines generated from a founder population of male and female N:NIH stock rats based on the capacity for energy transfer (i.e., aerobic capacity). Through an intricate breeding program and the selection for the polygenic trait of running capacity, HCRs/LCRs maintain genetic diversity but are differentiated by intrinsic mitochondrial function^23^. Because of this selective breeding over multiple generations, HCRs have greater mitochondrial content, oxidative capacity, and VO_2max_ compared to LCRs^23–25^. Further, LCRs develop diseases such as metabolic syndrome, fatty liver disease, and cancer ^23^ earlier and more frequently than HCRs and have a decreased lifespan^23,24^. Importantly, these differences emerge without exercise training, allowing for examining intrinsic mitochondrial function independently from exercise-training effects. Thus, the HCR/LCR model is an excellent model to test if differences in intrinsic aerobic capacity impact metformin treatment outcomes.

We aimed to test if outcomes from metformin treatment depend on intrinsic mitochondrial capacity. We used the HCR/LCR model because of the differing intrinsic mitochondrial oxidative capacity between the strains, and the genetic heterogeneity improves translatability to humans. We treated the rats at 18-months of age as it equates to middle-age when humans are likely to undergo healthspan-increasing treatment. We primarily focused on skeletal muscle because of its importance in maintaining metabolic control and mobility with age, but also examined the liver because of the effects of metformin on glucoregulation^26^. Our primary hypothesis was that mitochondrial remodeling to the energetic stress of metformin would differ between HCRs and LCRs. Thus, we hypothesized that metformin would result in positive mitochondrial remodeling in LCRs but would be either inconsequential or potentially detrimental for energetics in HCRs. Finally, we included a metformin washout group to help differentiate the long-term effects of metformin treatment from the acute presence of metformin in the tissue.

## Results

### Metformin dosing was similar among groups

HCR and LCR male rats were singly housed and randomly assigned to a control group (CON), a metformin-treated group (MET), or a metformin-treated group with a 48-hour washout of metformin prior to euthanasia (MET-WO). For the first seven days, the MET and MET-WO groups received 100 mg/kg/day of metformin in drinking water with non-caloric flavoring (Mio) to improve palatability (**Fig. 1**). After the first seven days, we increased the dose to 200 mg/kg/day for the remainder of the study. The MET-WO had metformin removed 48-hours prior to euthanasia. Control rats received tap water with non-caloric flavoring. Animal weight was measured weekly and water consumption was monitored daily to adjust metformin concentration in water to maintain the appropriate dose based on individual weight and water consumption of each animal (**Extended Data Fig. 1a,b**).

**Figure 1:**
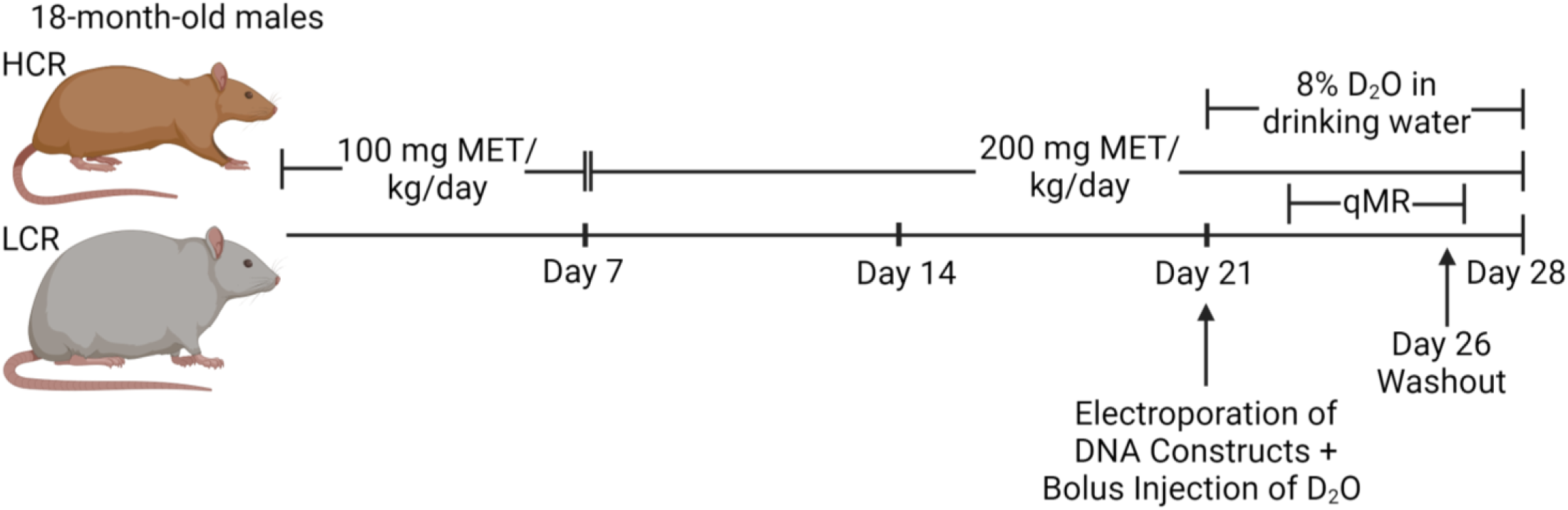
Study design. Age-matched, 18-month-old HCR and LCR male rats were randomly assigned to one of three groups: a control group (CON), an acute metformin group (MET), a 48-hour washout metformin group (MET-WO). The MET and MET-WO groups received 100 mg/kg/day of metformin in drinking water for the first 7 days of treatment. After day 7, the metformin dose was increased to 200 mg/kg/day. The MET rats received metformin until the day of sacrifice, while the MET-WO rats were switched back to tap water 48-hrs prior to euthanasia. The CON rats received regular tap water. All rats underwent deuterium oxide (D_2_O) stable isotope labeling during the last week of the intervention.

### Metformin induces a modest but positive effect on body composition in the LCRs with potential detriments to muscle mass in HCR

As expected, LCRs were heavier, had greater fat mass percentage, and lower lean mass percentage than the HCRs (main effect of strain, **Fig. 2a-2c**). Metformin affected the body composition of HCRs and LCRs differently. In LCRs, there were overall positive effects of MET where body mass was lower primarily due to a lower fat mass percentage with muscle masses (**Extended Data Fig. 1**). In contrast, the lower body mass in HCR-MET compared to HCR-CON appears to be due to loss of muscle mass (recorded in four of the five muscles) with no loss of fat mass. With MET-WO there were unexpected differences in body composition over such a short period of time. LCR-MET-WO had a greater fat mass percentage compared to the LCR-MET (**Fig. 2b**) and greater lean mass percentage compared to CON (**Fig. 2c**), while the HCR-MET-WO had a greater lean mass percentage compared to CON and MET (**Fig. 2c**). Because of the unexpected differences in body composition from acute MET-WO, we analyzed body water percentages and found that differences in body water percentage tracked with differences in percent lean mass (**Fig. 2i**). We measured gastrocnemius (gastroc) glycogen concentrations to determine if the shifts in body water were from differences in muscle glycogen storage (**Fig. 2j**). There were notable differences in muscle glycogen storage between the stains, and significantly lower glycogen concentrations after MET-WO in the HCR (but not LCR). However, these differences could not explain the changes in the body water pool. One further insight into the potential changes in the body water pool comes from our labeling with deuterium oxide (D_2_O) (to measure protein synthesis) over the last week of the study. In both strains, the MET-WO has higher body water enrichment than MET (**Extended Data Fig. 1**). When consuming D_2_O-supplemented drinking water, there can be differing resultant body water pool deuterium enrichments between animals because of differences in metabolically produced water. Our data show that potential changes in macronutrient oxidation (potentially greater fat oxidation) during MET-WO impacted the body water pool size, confounding measurements of body composition. We conclude that there were benefits to body composition in LCRs, but potential detriments to muscle mass in HCRs.

**Figure 2:**
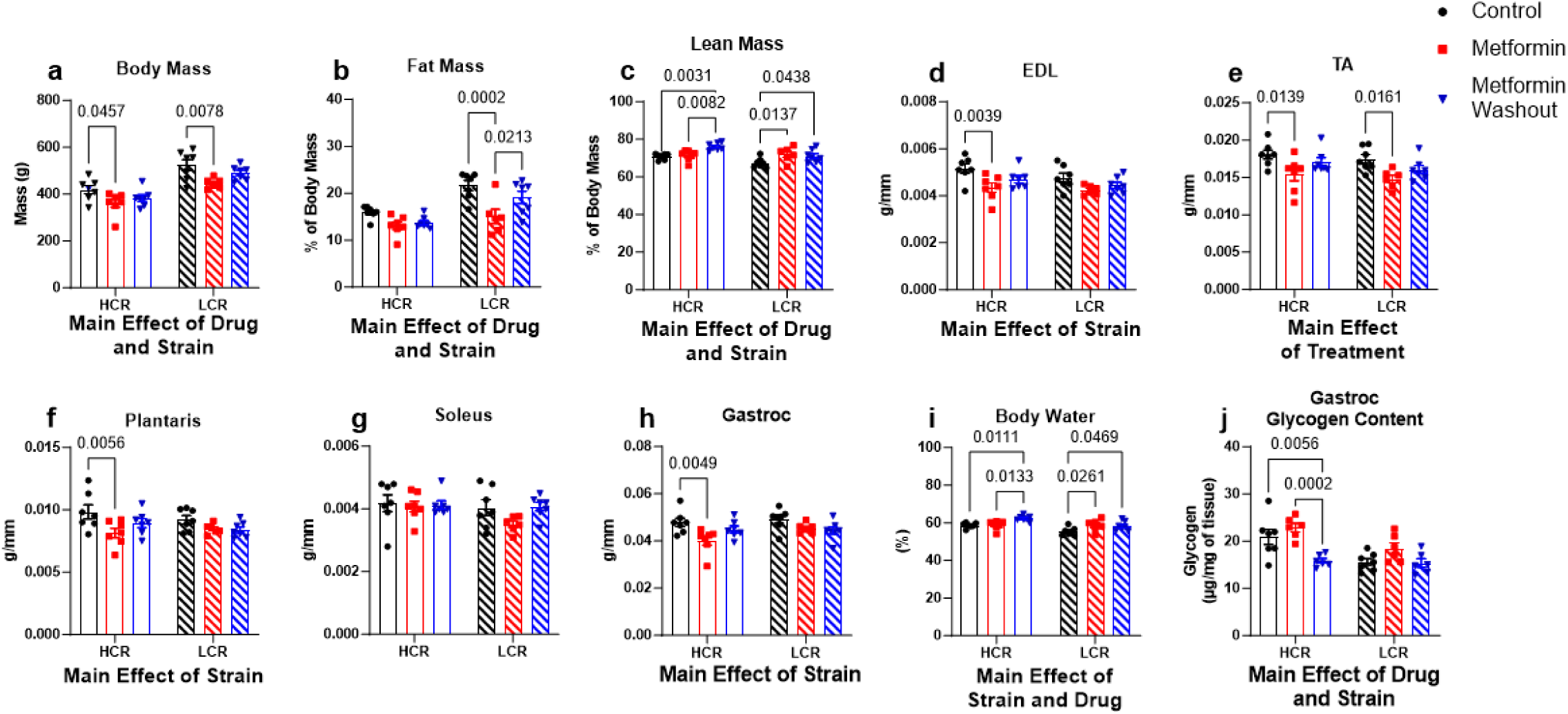
**a-c,** Body composition measures from CON (black), MET (red) and MET-WO (blue). **d-h**, Skeletal muscle masses normalized to tibial length. **i**, Total body water percentage. **j**, Gastroc glycogen content. Data were analyzed by two-way ANOVA (group x treatment) and are represented as mean ± SEM from 6-7 rats per group.

### AMPK is elevated in response to the removal of metformin

Previous studies have indicated that the primary mechanism of action of metformin is through CI inhibition or reducing mitochondrial membrane potential^14,15,17^. The resultant energetic stress activates AMPK, which initiates a host of downstream effects^27^. In HCRs, after four weeks of MET there was no difference in AMPK activation in muscle (**Fig. 3**) or liver (**Extended Data Fig. 2c**) compared to CON. This finding indicates that MET did not activate AMPK in HCRs, or that compensatory changes over the four weeks corrected any energetic stress. However, in skeletal muscle (but not liver) of HCRs, 48-hours of MET-WO resulted in higher levels of AMPK phosphorylation compared to CON or MET, indicating that the MET-WO created an energetic stress. In LCRs, MET resulted in higher AMPK phospho/total ratios compared to CON, but this was primarily due to lower total AMPK rather than higher phosphorylation. With MET-WO, this difference from CON remained without any further changes from the MET (**Fig. 3**). Although the primary site of the therapeutic benefits of MET are thought to be at the liver, we found little evidence that MET differentially impacted HCRs and LCRs at the liver as evidenced by no changes in liver mass, triglycerides, or transcription of the gluconeogenic enzymes PEPCK or G6Pase (**Extended Data Fig. 2**).

**Figure 3:**
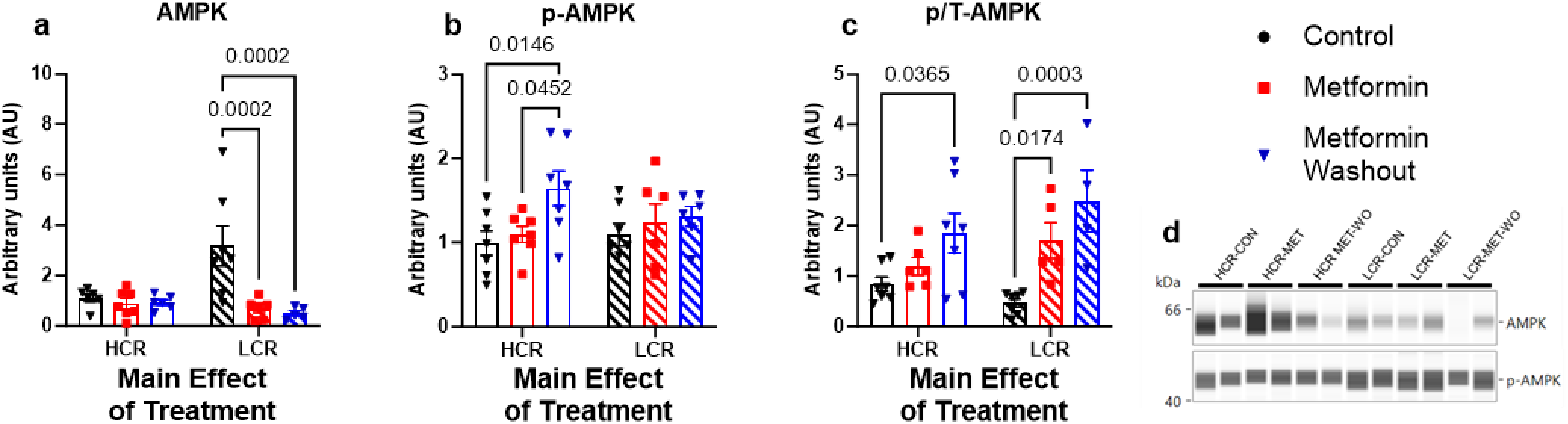
Determination of the nutrient sensing marker AMPK, **a**, total, **b**, phosphorylated, and **c**, phospho:total ratio in the gastroc. Data were analyzed by two-way ANOVA (group x treatment) and are represented as mean± SEM from 6-7 rats per group.

### HCRs and LCRs have heterogenous mitochondrial remodeling in response to metformin

Protein remodeling is a primary cellular mechanism of stress adaptation. Protein synthesis is the most energetically costly cellular process^28^, which can be compromised during periods of energetic stress. We therefore performed a series of analyses to determine if MET impaired protein remodeling in skeletal muscle, and if the adaptations of mitochondrial proteins to MET differed between the strains. By providing D_2_O in drinking water over the final week of treatment, we measured the cumulative protein synthesis over that period when a new steady-state was assumed to be achieved. As a general indicator of whether a potential energetic stress from MET compromised protein turnover, we investigated bulk protein turnover in the gastroc, soleus, and tibialis anterior (TA) (**Fig. 4**) and liver (**Extended Data Fig. 2g,h**). There were no differences in bulk protein turnover of any tissue protein fraction with MET indicating that protein turnover was not compromised.

**Figure 4:**
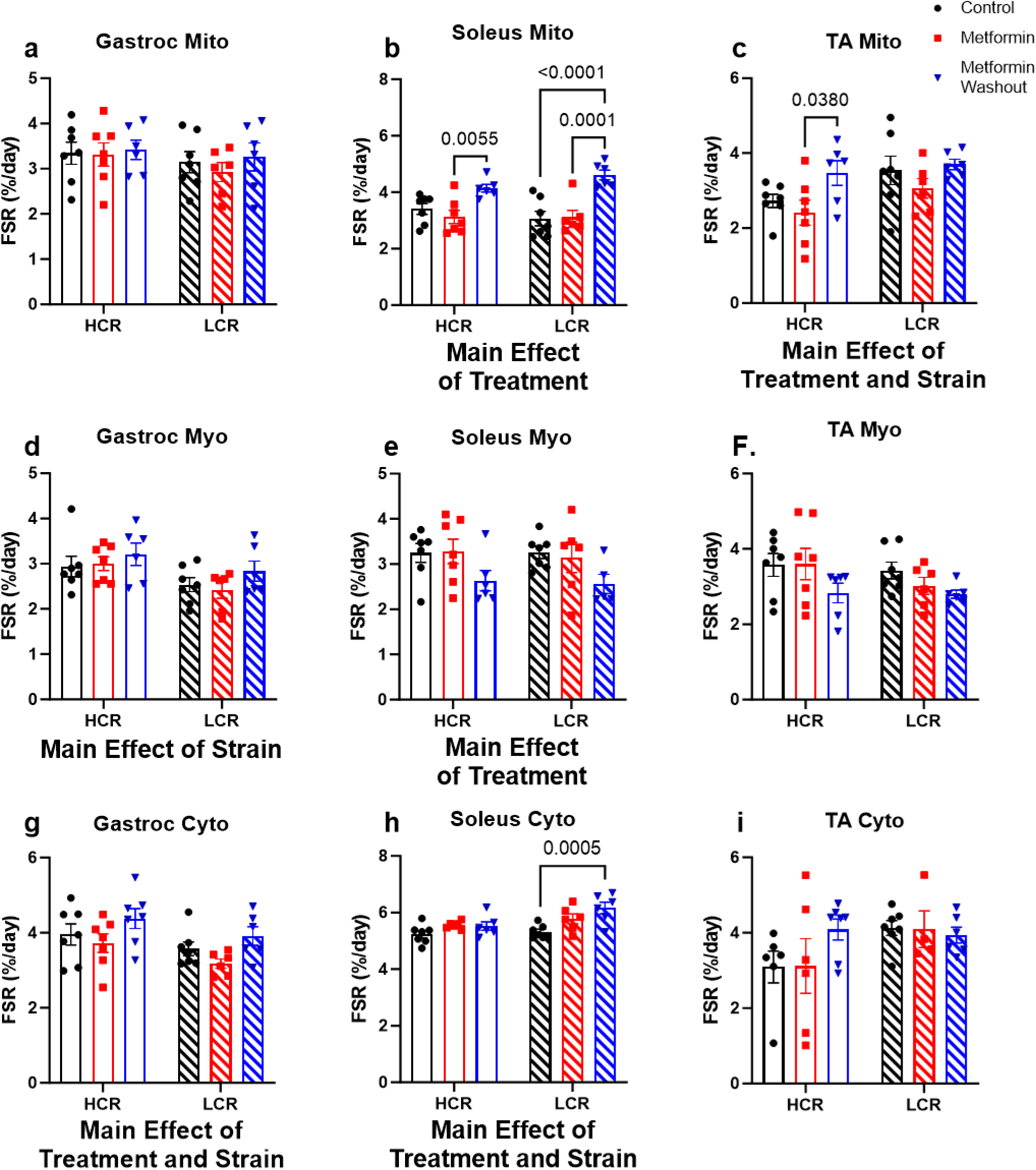
Determination of the fractional synthesis rates in **a-c**, mitochondrial (Mito), **d-f**, myofibrillar (Myo) fractions, and **g-I**, cytosolic (Cyto) of the gastrocnemius (gastroc), soleus, and tibialis anterior (TA) muscles. Data were analyzed by two-way ANOVA (group x treatment) and are represented as mean ± SEM from 4-7 rats per group.

Mitochondria are one of the primary targets of metformin^14,15,17^. Therefore, we used a targeted proteomic analysis to examine changes to protein concentrations of individual proteins in the ETS of gastroc and soleus^29^. We used the gastroc and soleus due to their different fiber type compositions and mitochondrial content. Our targeted panel consisted of 62 proteins; 28 from CI, three from Complex II (CII), 12 from Complex III (CIII), eight from Complex IV (CIV), and 11 from Complex V (CV) (details of validation are provided **Supplemental Table 1**). We present our data as dot plots to visualize concentrations relative to each other (**Fig. 5**) and with 3D visualization to determine if there was a spatial relationship (e.g., within or between complexes) of any changes in concentration (**Extended Data Fig. 3 and 4;** video representations are available at DIO:10.6084/m9.figshare.25301230). In the gastroc and soleus, there was a main effect of strain where HCRs had greater concentration of ETS proteins compared to LCRs (**Fig. 5a,b**). After four weeks of MET in HCRs, there were no differences in the concentrations of any ETS proteins in the gastroc when compared to CON (**Fig. 5a**). However, in the soleus, HCR-MET had lower concentrations of 18 (six CI, three CIII, four CIV, and five CV) proteins compared to the CON. Half of the proteins were in the mitochondrial matrix, while the other half were in the mitochondrial membrane (**Extended Data Fig. 4**), suggesting no spatial relationship. In contrast to HCRs, MET did not change mitochondrial protein concentrations in either gastroc or soleus of LCRs (**Fig. 5b and Extended Data Fig. 4** and 5). The measurement of proteins concentrations provides insight into net changes, but not dynamic remodeling since protein turnover can increase without changes in concentration.

**Figure 5:**
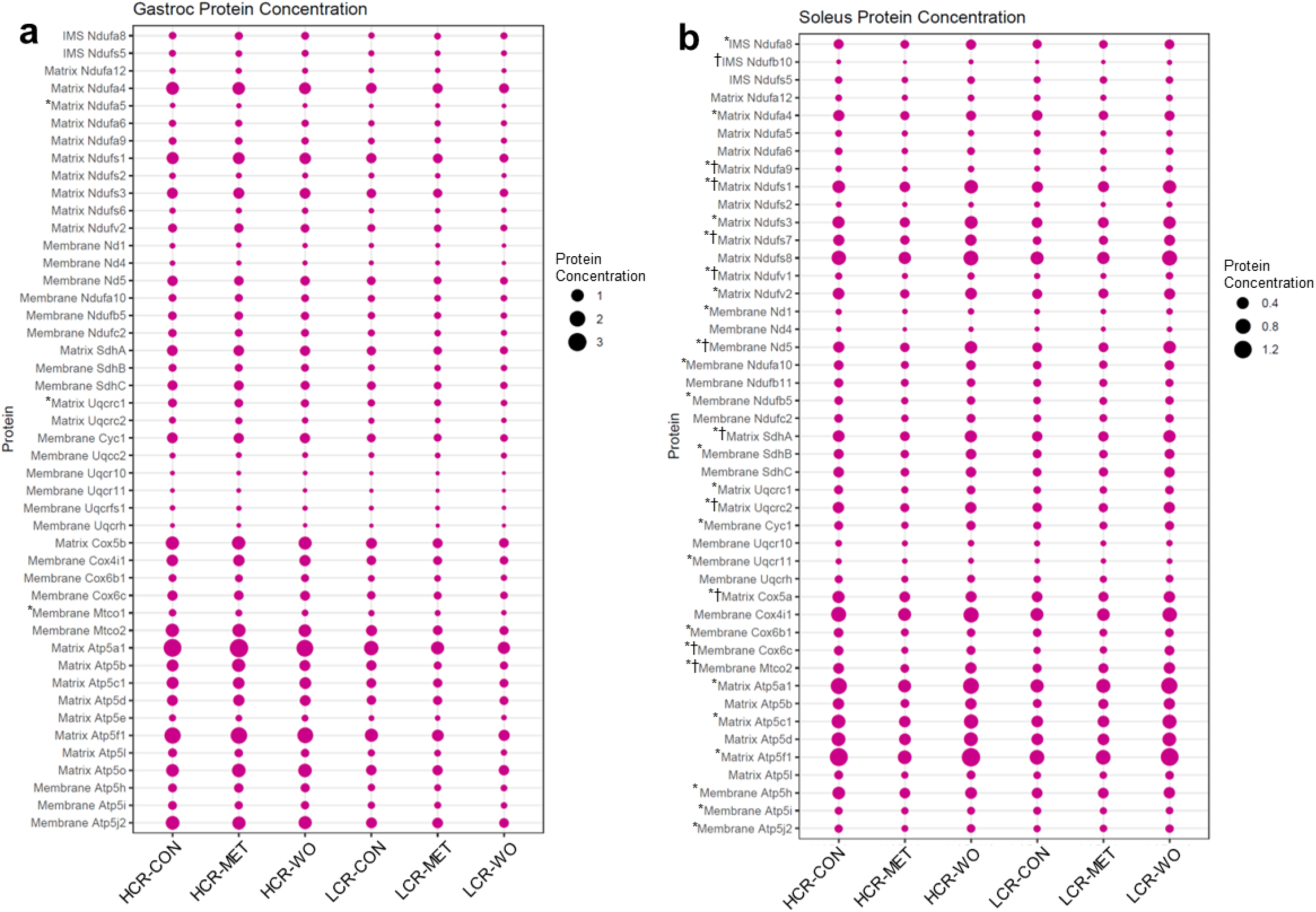
We performed a mitochondrial targeted quantitative proteomics in rat **a**, gastroc and **b**, soleus skeletal muscles. The size of the dot indicates the relative abundance of the protein. Data were analyzed by two-way ANOVA (group x treatment) and are represented as mean ± SEM from 5-7 rats per group. * indicates significant difference within HCRs and † indicates significant difference within LCRs.

To determine the dynamic remodeling of the mitochondrial proteome, we further analyzed the proteomic data for the synthesis rates of the individual proteins by the incorporation of D_2_O. By applying the three-dimensional modeling, we observed that in general proteins in the mitochondrial membrane for CI and CV (but not CII, CIII, or CIV) turnover at a slower rate compared to those outside to the mitochondrial membrane (**Fig. 6a,b**), which has been previously observed^30^. In the gastroc of the HCRs, MET had slower mitochondrial protein synthesis rates for CI (despite no changes in concentration) compared to CON, while other complexes were largely unaffected (**Fig. 6a**). In the soleus of HCRs, 11 proteins changed synthesis rates (6 proteins had higher synthesis rates: two CI, one CIII, one CIV, and two CV and 5, while five proteins had lower synthesis rates: three CI, two CIII, and one CIV) in MET compared to CON (**Fig. 6b**). In the soleus of the LCRs, MET resulted in 13 CI proteins having greater synthesis rates, of which four proteins are core subunits, and one protein had a slower synthesis rate compared to CON. Also, in LCRs, seven of 10 CV proteins (6/7 proteins were in the matrix) had greater synthesis rates compared to CONs, while there was little impact on CV in HCRs (**Fig. 6b**). It is notable that changes in turnover of the mitochondrial proteins did not always match with changes in concentration. Collectively, our quantitative and kinetic proteomics suggest that metformin causes more mitochondrial remodeling in LCRs than HCRs.

**Figure 6:**
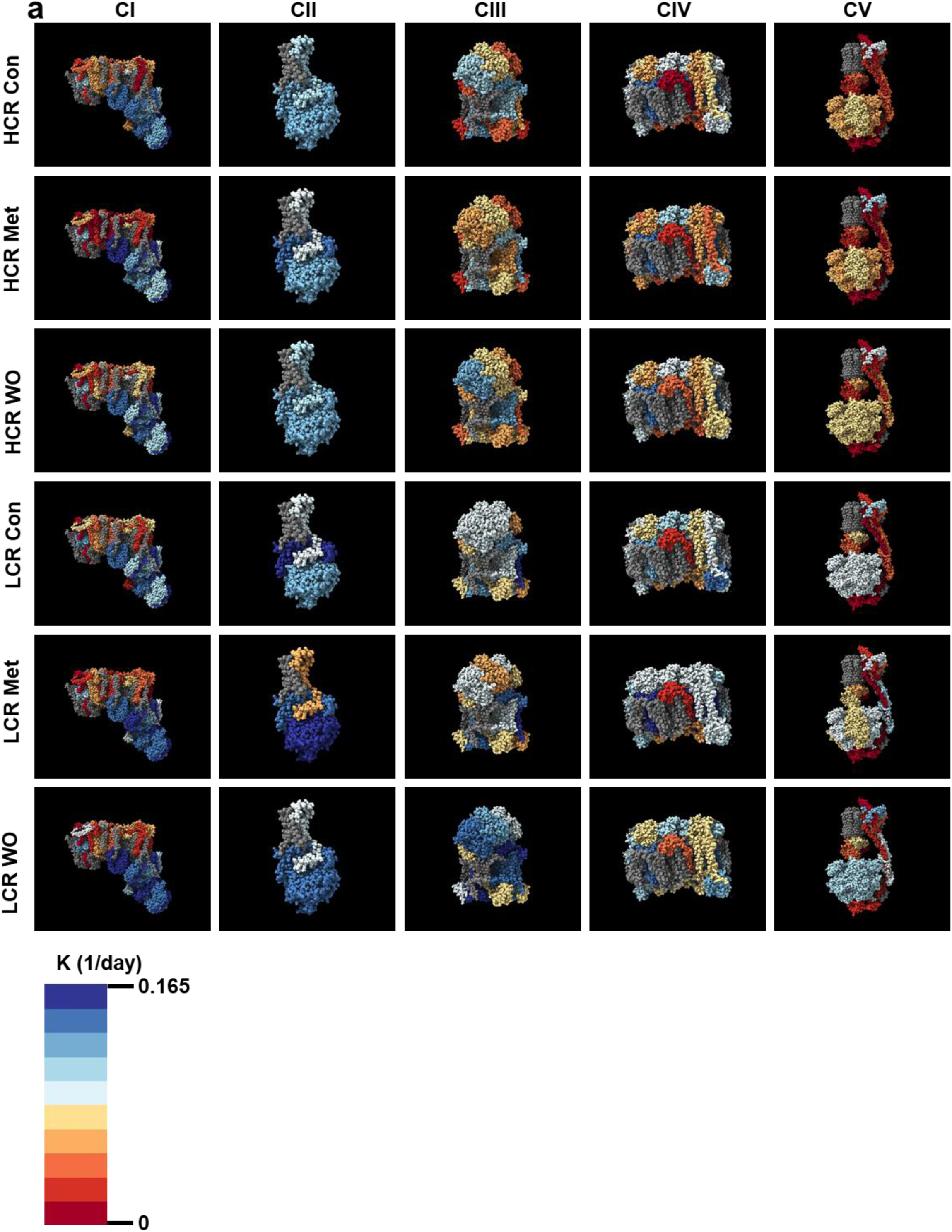
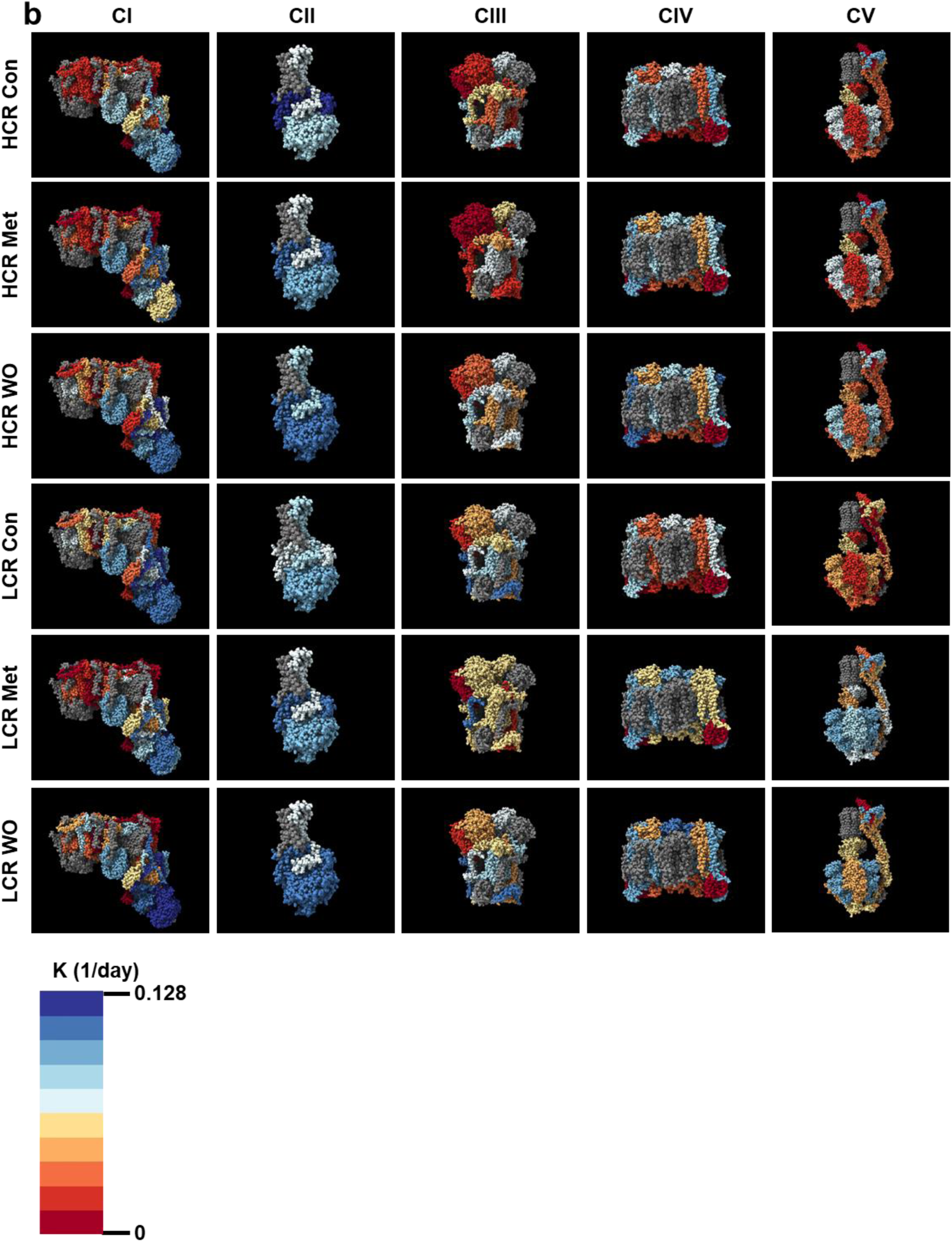
Visualization of the fractional synthesis rates of the mitochondrial ETS complexes in the **a**, gastroc and **b**, soleus. Gray proteins indicate that the protein was not detected in our analyses. The complexes were generated using complex IDs (CI: 7AK5, CII: 1ZOY, CIII: 7TZ6, CIV: 7COH, and CV: 8H9V) from the RCSB Protein Data Bank and rendered using ChimeraX. The data are represented as mean from 5-7 rats per group.

### Metformin removal induces rapid mitochondrial remodeling that is distinct in each strain

When we removed metformin, there were surprising changes in abundance and synthesis in that short period of time. In the gastroc of the HCRs, where MET had minimal differences in turnover, there were substantial differences in synthesis rates among the complexes (**Fig. 6a**). In HCR-MET-WO, most of the proteins were greater than MET (similar to CON), while a few proteins were greater than the CON, suggesting a rebound in synthesis rates after the WO. In the gastroc of LCRs, MET resulted in greater synthesis rates compared to CON, while the MET-WO resulted in greater synthesis rates compared to MET and CON (**Fig 6a**). These changes in synthesis rates resulted in minor changes in mitochondrial protein concentrations (**Fig. 5**). In contrast to the gastroc, the changes in synthesis rates of the individual mitochondrial proteins from the MET-WO resulted in changes in protein concentrations in the soleus (**Fig. 5b and 6b**). These changes in synthesis and concentration were evident in both strains, although the impacted proteins were not always the same between the strains (**Fig. 6b**). In the case of the LCRs, the MET-WO made the mitochondrial proteome concentrations similar to HCR-CON (**Fig. 5b**). It is worth reiterating our labeling approach to grasp the magnitude of the changes in synthesis over the short MET-WO period. Our labeling approach measures cumulative synthesis of proteins over the last week. The MET-WO was only the last 48-hours of that two-week period, so the degree of the changes apparent with MET-WO are above the similar amount of synthesis that would have occurred in MET and MET-WO groups up to that point. Lastly, it is important to note that there were exceptions to these patterns within the MET-WO of both strains, suggesting some level of heterogeneity in the response to MET-WO between strains.

### Removal of metformin induces mitochondrial fission in the subsarcolemmal population

The second primary mechanism of mitochondrial remodeling are fission-fusion events that change mitochondrial morphology^31^. We first performed western blotting on protein markers of fission and fusion. Although there was a main effect of strain on MFF, MFN2, and OPA1 (**Fig. 7a,d,e**), there were no differences within a strain from MET or MET-WO (**Fig 7a-e**). Given the limited insight provided by these snapshots, we examined changes in mitochondrial morphology. To do so, we electroporated the TA with constructs to label mitochondria with mitochondria-targeted fluorescent reporters. We used TA due to its relative ease for electroporation. Skeletal muscle mitochondria reside in two spatially distinct subpopulations: subsarcolemmal mitochondria (SS) found along the cell perimeter and interfibrillar mitochondria (IFM) located between the myofibrils. These populations are functionally distinct and respond to stimuli differently^32,33^. To our knowledge, there is no data on the effects of metformin on these mitochondrial populations in skeletal muscle. For our analyses, SS were defined as the mitochondria located in the outer 10% of the cross section of the fiber. MET did not change mitochondrial morphology in HCRs (**Fig. 7f-n**) but resulted in a higher ratio of mitochondrial CSA to muscle fiber CSA in LCRs (**Fig. 7k**). However, 48-hours of MET-WO caused differences in the distribution of mitochondria in both strains that were largely driven by decreased SS area (**Fig. 7g**). In both strains, compared to MET, MET-WO had lower SS area (**Fig. 7g**) and SS CSA/total mitochondrial CSA (**Fig. 7i**), suggesting that the MET-WO induces rapid changes to the SS population. These results were somewhat expected as the SS population responds more rapidly to stimuli and are less spatially restricted than IMF ^32,34^. When expressed relative to muscle fiber CSA, IMF CSA was higher after MET-WO in HCRs, but lower in LCRs (**Fig. 7m**). In sum, by four weeks of MET, the mitochondrial pool was expanded relative to fiber CSA in LCR, but no changes in HCRs. However, MET-WO rapidly changed morphology with a shift toward greater IMF relative to muscle CSA in HCRs but lower IMF in LCRs. Our results clearly demonstrate differences in the number, size, and percentage of SS in HCRs vs LCRs TA muscle fibers, and that metformin impact these mitochondrial populations differently.

**Figure 7:**
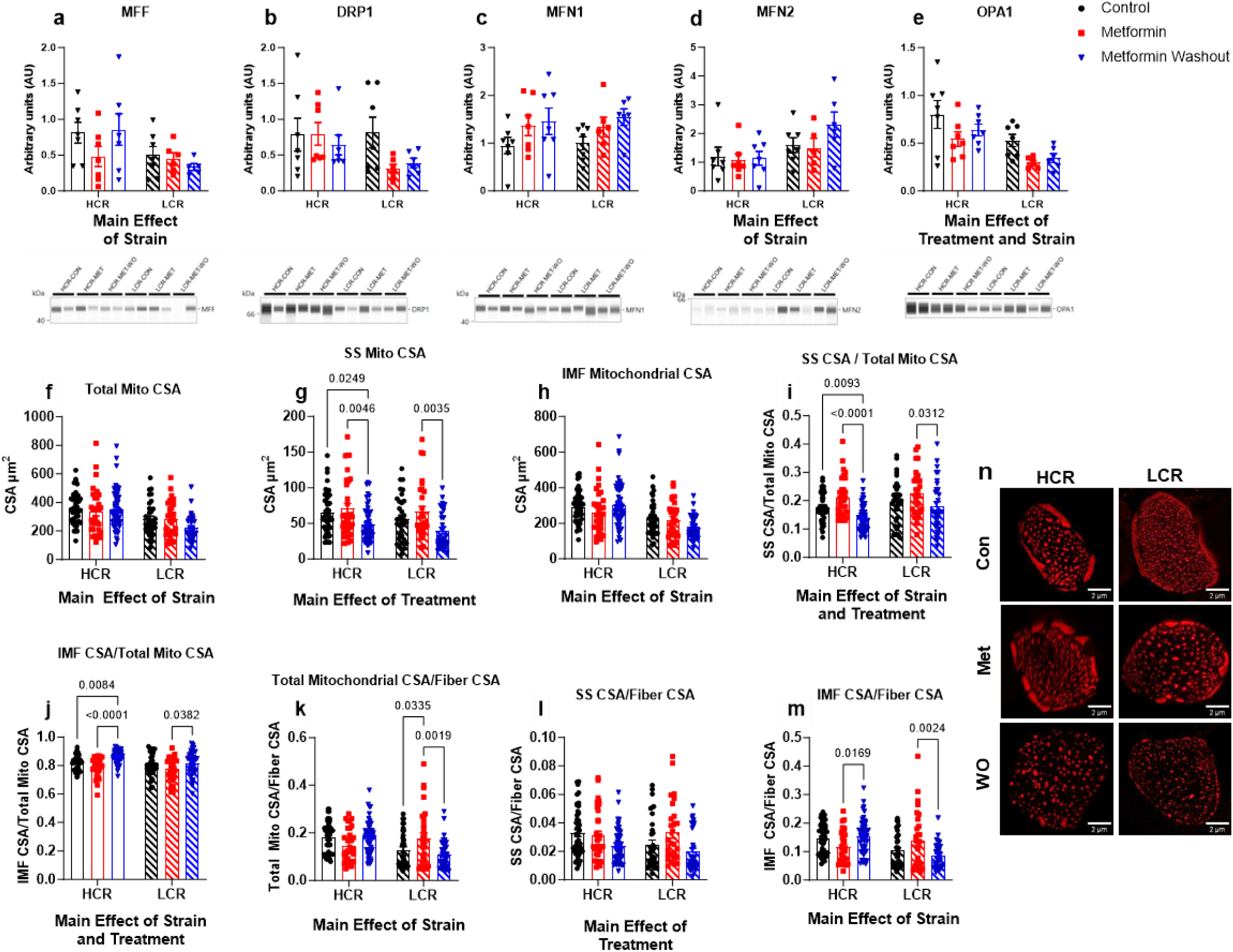
**a-e**, Mitochondrial dynamics markers. The average CSA of the **f**, total mitochondria, **g,** subsarcolemmal population, and **h,** intermyofibrillar population. **i**, The total subsarcolemmal CSA normalized to total mitochondrial CSA. **j**, The total intermyofibrillar CSA normalized to total mitochondrial CSA. **k**, The total mitochondrial CSA normalized to total fiber CSA. **l**, The total subsarcolemmal CSA normalized to total fiber CSA. **m**, The total intermyofibrillar CSA normalized to total fiber CSA. **n**, Representative images. Data were analyzed by two-way ANOVA (group x treatment) and are represented as mean ± SEM from 3-7 fibers taken from a single image from 6-7 rats per group.

### Metformin inhibits mitochondrial respiration, but the washout increases respiration capacity in HCRs

To assess the resultant differences in mitochondrial respiratory function from the remodeling period, we used high resolution respirometry. We used our established ADP titration protocol that previously showed that MET inhibited the positive effects of exercise^12^. This protocol uses CI substrates and an ADP titration to determine ADP sensitivity^12^ and maximal oxygen consumption. After establishing maximal respiration (V_max_), we use succinate to determine maximal CI+II activity, and rotenone to inhibit CI to determine effects downstream of CI.

As expected, there was a main effect of strain where HCRs had a higher V_max_ than LCRs (**Fig. 8a**) demonstrating the difference in intrinsic aerobic capacity between HCR and LCRs. Four weeks of MET impaired respiration at higher ADP concentrations in LCRs and HCRs, although in HCRs this impairment displayed a trend (p=0.06, **Fig. 8b,c**). These data show that our proposed dose of metformin, which is equal to clinical doses in humans and the TAME trial^4^, impairs respiration in HCRs and LCRs (up to 40% and 50%, respectively). In rats with a 48-hour MET-WO, there was a striking change in respiration that differed by strain. In HCRs, MET-WO increased O_2_ flux at all concentrations of ADP to a level that exceeded both CON and MET. In LCRs, the respiration rate in MET-WO was greater than LCR-MET, but was equal to CON. To test whether potential differences were the result of the presence or absence of MET, we used fiber bundles from the MET-WO groups in a separate instrument and spiked the respiration chamber with MET. The metformin-spike had no impact on O_2_ flux in LCRs, but slightly impaired respiration in HCRs where O_2_ flux was still significantly higher than MET, but no longer different than CON. These differences with MET-WO in HCRs persisted when succinate was added to the chamber and when CI was inhibited with rotenone (**Extended Data Fig. 5**), indicating that the increase in O_2_ flux during MET-WO was not solely a CI effect. Further, these differences were apparent when uncoupling. We therefore looked at the oxphos to electron transport capacity ratio (P/E) as an indicator of potential deficiencies in oxphos (**Fig. 8d**). There were no differences in the P/E with any treatment in either strain, indicating that impairments and compensatory changes in mitochondria were due to the effect of MET on electron transfer rather than phosphorylation. In sum, it appears that during metformin treatment, HCRs make better compensatory changes to maintain respiration compared to LCRs. These compensatory changes become apparent in HCRs when metformin is removed, and flux is much greater compared to CON or MET, but in LCRs the changes are not different from CON. Since the MET-Spike changes flux to not be different from CON in HCRs, but does not further impact LCRs, we conclude that the acute presence of MET differentially impacts O_2_ flux in rats with high or low intrinsic aerobic mitochondrial function (i.e., context specific).

**Figure 8:**
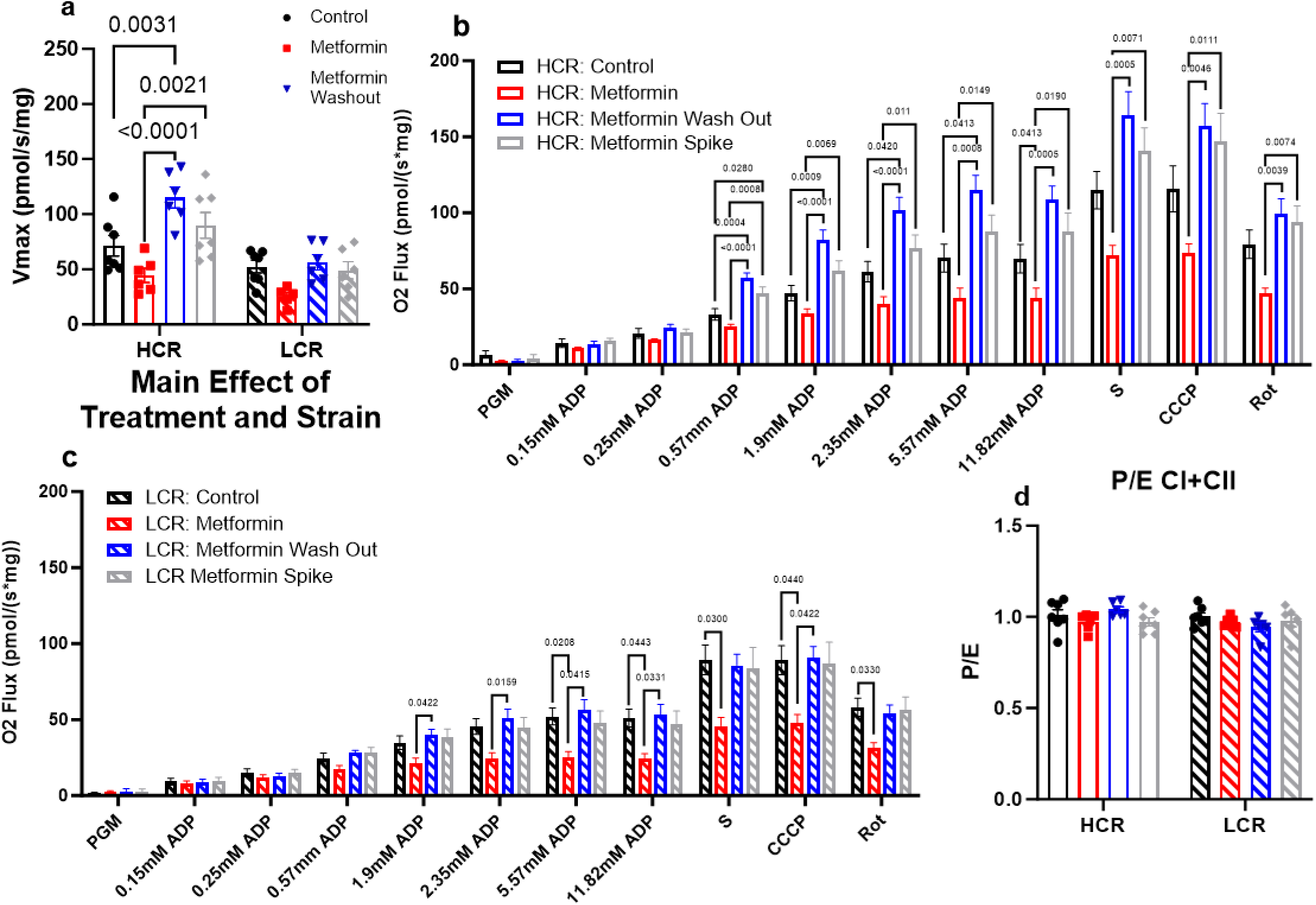
**a**, Maximal mitochondrial respiration (V_max_). Mitochondrial respiration in **b**, HCRs and **c**, LCRs. The apparent Km of ADP was calculated using Michaelis-Menten Kinetics. Data were analyzed by two-way ANOVA (**a** and **d**) or by one-way ANOVA for each substrate addition (**b** and **c**) within each strain. Data are represented as mean ± SEM from 4-7 rats per group.

## Discussion

This study shows complex changes in muscle mitochondrial structure and function with four weeks of metformin treatment that changed rapidly with the removal of metformin. These changes to metformin treatment and its subsequent removal were different depending on intrinsic mitochondrial function (HCRs vs LCRs), the oxidative capacity of the muscle (gastroc vs soleus), and the mitochondrial population (IMF vs SS). Metformin caused lower ADP-stimulated respiration in LCRs, with less of a change in HCRs. However, the washout of metformin resulted in a doubling of respiratory capacity in HCRs, but more modest increases in LCRs. These improvements in respiratory capacity appeared to be caused by mitochondrial remodeling that included increases in protein synthesis and changes in morphology. In line with our hypotheses, the impact of metformin differed by intrinsic mitochondrial function. However, the changes were more complex than anticipated and dependent on whether metformin was acutely present. From this relatively short treatment, there is context specificity to metformin treatment, although how these effects manifest over the long-term remain unclear.

Metformin treatment had positive effects on body composition in the LCRs, with an unexpected loss of muscle mass in HCRs. Prior reports have shown that the quadriceps and soleus muscles of db/db mice decrease in response to metformin treatment while the EDL, gastroc, and TA were unaffected suggesting the glucose-lowering effect of metformin maybe partially offset by loss of muscle mass^35^. These changes in muscle mass likely have functional consequences. Changes to skeletal muscle mitochondrial structure and function were strain specific. Our measurements of mitochondrial respiration and morphology were after four-weeks of metformin treatment and mitochondrial protein turnover were cumulative changes over the last week. Therefore, it is hard to say that the observed remodeling caused changes in mitochondrial respiration or that changes in mitochondrial respiration triggered the remodeling. However, after metformin treatment the concentration of several mitochondrial proteins was lower in the soleus of HCRs compared to controls. This same decrease was not observed in LCRs and there was in fact a greater mitochondrial CSA. In both strains, the changes in mitochondrial protein concentrations were more prominent in the oxidative soleus compared to the more mixed gastroc. There were changes in synthesis rates of the individual mitochondrial proteins although these differed by strain. Despite these changes, LCR+MET had significantly lower mitochondrial respiration than CON, while the differences in HCR did not reach significance.

The removal of metformin provided additional insight into how metformin impacts skeletal muscle outcomes under different contexts. After 48-hours of metformin washout, cumulative synthesis rates of mitochondrial proteins were greater, with individual proteins driving the effect. These changes were accompanied by increased mitochondrial protein concentrations, and a shift of mitochondrial area from SS to IMF which together presumably drive the robust increase in mitochondrial respiratory capacity. Our interpretation is that during the metformin treatment HCRs are largely able to maintain mitochondrial respiratory function. However, removal of the metformin triggers remodeling with a robust rebound (and overshoot) of aerobic capacity. The observed rapid remodeling likely occurs because of the release of a metformin-induced brake produced by the accumulation of positively charged metformin in the mitochondrial matrix. These findings could explain our previous study where we found that metformin hinders mitochondrial adaptation to exercise^12^. It is possible that although the muscle mitochondria can manage to adapt to metformin on board, it cannot further adapt to the addition of an energetic stress. The fission of the SS, as suggested by lower SS CSA relative to total mitochondrial CSA, in response to the washout is likely a contributing factor to the greater mitochondrial respiration capacity in HCRs. The SS turns over at a faster rate and alters oxidative capacity to a greater extent than the IMF in response to changes in energetic demand^36–39^. Although we see similar SS mitochondrial morphology in LCRs compared to HCRs, we did not observe any changes in mitochondrial respiration in LCRs after MET-WO. This failure to see the greater mitochondrial respiration in LCR-MET-WO may be due to a shift in cellular priorities.

There are a few limitations to the current study. First, we did not assess changes that occurred early in response to metformin. There likely were changes in protein synthesis earlier in the treatment phase that we did not capture. Although we were interested in change during chronic metformin treatment, early changes from metformin would have been interesting to assess from a remodeling perspective. Second, it could be argued that four weeks of treatment is not long-term; thus, we cannot determine changes in healthspan related outcomes. We are concerned with the long-term healthspan extension applications of metformin treatment; therefore, our window of application is limited as there is 100% survival at 16 months for both strains, which begins to decline at roughly 18 months for LCRs^40^. Thus, our metformin treatment begins about at the timepoint when healthspan is in decline. Third, we did not conduct mitochondrial respiration assays in the soleus or TA. The soleus and TA showed large, distinct changes in our proteomic and morphology analyses, respectively. These data could help elucidate the fiber type differences see with metformin. However, this was not feasible due to the low throughput of high-resolution respirometry. Finally, these studies were performed in male rats as this was a preliminary study into the effects of metformin in rats with different intrinsic aerobic capacity. Thus, we cannot make conclusions about the context-specific effects of metformin in female rats or between sexes. Future studies should further delineate the timing of changes in oxidative capacity and mitochondrial remodeling to determine causation and add energetic challenges such as exercise to determine limits to remodeling.

## Conclusion

The differences in strains of rats with distinct intrinsic mitochondrial aerobic capacities draw into question whether the positive findings of metformin treatment in human or model organisms that are less metabolically healthy can be broadly applied to all metabolic conditions^4^. We raise this concern because treatments which prolong healthspan would logically start while an individual is healthy and absent of chronic disease^9,41^. Our current study demonstrates that in skeletal muscle, which is crucial for metabolic health, mobility and mortality, there are clear differences in the effects of metformin based on background metabolic health. Additionally, we show that the observed changes from metformin treatment can be dramatically different depending on the acute presence or absence of metformin since there can be (in the appropriate context) rapid mitochondrial remodeling after removal. The findings from this study are critical given the ever expanding off-target use of metformin in healthy individuals without chronic disease and/or overt metabolic dysfunction.

## Methods

### Ethical approval

We performed all animal procedures in accordance with protocols approved by the Institutional Animal Care and Use Committee at the Oklahoma Medical Research Foundation and the guidelines provided by the National Research Council’s Guide for the Care and Use of Laboratory Animals: Eighth Edition.

### Animals

We obtained male, 18-month-old high- and low-capacity runners (HCR/LCR) of the 44^th^ and 45^th^ generation from The University of Toledo. The rats were bred and maintained as previously described^24^. Upon arrival, the rats were randomly assigned to a group and housed individually in a temperature-controlled environment with a 12:12-hr light-dark cycle. Throughout the study, rats were given *ad libitum* access to food (PicoLab Rodent Diet 20, cat #: 3005750-220, 62.4% CHO 24.5% PRO, and 13.1% FAT) and water.

### Experimental design

HCR and LCR male rats were singly housed and randomly assigned to one of three groups: a control group (CON), a metformin-treated group (MET), and a metformin-treated group with a 48-hour washout of metformin prior to euthanasia (MET-WO). For the first seven days of treatment, the MET and MET-WO groups received 100 mg/kg/day of metformin in drinking water with non-caloric flavoring (Mio) to improve palatability (**Fig. 1**). After the first seven days, we increased the dose to 200 mg/kg/day for the remainder of the study. The MET-WO had metformin removed 48 hours prior to sacrifice. Control rats received regular tap water with non-caloric flavoring throughout the experiment. Animal weight was measured weekly, and water consumption was monitored daily throughout the treatment to adjust metformin concentration in water to maintain the appropriate dose based on the individual weight and water consumption of each animal. On day 21, all rats received an IP bolus injection of filtered 99% D_2_O (Sigma Aldrich, St. Louis, MO) equivalent to 5% of body water (estimated at 60% of body mass), followed by 8% D_2_O-enriched drinking water for the remaining seven days. Additionally, on day 21 we electroporated one TA with constructs to visualize the outer mitochondrial membrane and matrix. Body composition was measured using quantitative magnetic resonance (QMR, Echo Medical Systems, TX, USA) during the terminal five days. Food was removed 12 hours prior to sacrifice. Blood from cardiac puncture was collected into EDTA-lined tubes on ice, and plasma was frozen. Tissues were removed, weighed, flash frozen, and stored at −80°C unless otherwise noted. The TA was fixed in 4% paraformaldehyde and cryo-embedded.

### Glycogen assay

We quantified glycogen content of the gastrocnemius (gastroc) according to previously established protocols^42^ using 20 mg of skeletal muscle powered in liquid nitrogen using 20 mg of skeletal muscle powered in liquid nitrogen. Approximately 20 mg of skeletal muscle was powered in liquid nitrogen and immediately placed in 200 µL of 0.5 M NaOH, which was then placed in a 100°C heating bath for 30 minutes. Then 50 µL of sodium sulfate and 600 µL of ethanol was added to each solution and centrifuged at 2000 RCF for 10 minutes. The resulting pellet and glycogen standards were resuspended in 500 µL of DI water and 300 µL of 95% sulfuric acid was quickly added followed by 45 µL of the 5% (w/v) phenol solution. The samples and standers were incubated for 30 minutes at room temperature. Lastly, 280 µL of reaction mixture to a microplate, and read in the plate reader at 488 nm in duplicate.

### Semiquantitative immunoblots

For all skeletal muscle data, approximately 20–30 mg of gastroc muscle was lysed and the protein concentration was determined by Peirce 660 assay (cat#: 22660, ThermoFischer Scientific, Walktham, MA) as described previously^43^. Lysed protein fractions were processed to visualize proteins by the ProteinSimple (San Jose, CA, USA) WES system. For primary antibodies we used AMPK (1:50, #2603), p-AMPK (1:25, #2603), Mitofusin-1 (1:50, #14739), Mitofusin-2 (1:50, #11925), MFF (1:50, #84580), DRP1 (1:50, #8570), p-DRP1 (1:50, #6319), OPA1 (1:50, #80471), and Tom20 (1:50, #42406 from Cell Signaling (Danvers, MA, USA). Secondary antibodies were included in a WES Master Kit (DM-001, ProteinSimple). Data were normalized to HCR-CON. Compass files can be found at DIO:10.6084/m9.figshare.25301230.

For liver analysis, tissue was homogenized in Cell Lysis Buffer (Cat#9806S, Cell Signaling, Danvers, MA, USA) with protease & phosphatase inhibitors (Cat#5872S, Cell Signaling, Danvers, MA, USA). We quantified total protein using BCA Protein Assay Regent Kit (cat#23227, Pierce, Rockford, IL, USA). Then we separated proteins into 4-15% gradient SDS-PAGE gels and transferred them to PVDF immuno-blot membranes (Bio-Rad, Hercules, CA). The primary antibodies S6 (1:1000, #2217S), p-S6 (1:1000, #2211S), AMPK (1:1000, #2532S), and p-AMPK (1:1000, #2535S, Cell Signaling, Danvers, MA, USA) were used according to manufacture instructions and IRDye 800CW Infrared secondary antibody (Cat #926-32213, LI-COR Biotechnology, Lincoln, NE) was used at a concentration of 1:15000. Imaging was done on an Odyssey Fc Imaging System (LI-COR Biotechnology).

### Liver triglycerides

We homogenized 100mg of liver in Cell Signaling Lysis Buffer (Cat#9806S, Cell Signaling, Danvers, MA) with phosphatase and protease inhibitors (Cat#5872S, Cell Signaling, Danvers, MA, USA). Total lipid was extracted from this homogenate using the Folch method with a 2:1 chloroform-methanol mixture as previously described^44^. A nitrogen drier at room temperature was used to dry the lipids prior to reconstitution in 100uL of 3:1:1 tert-butyl alcohol-methanol-Triton X-100 solution. Triglyceride concentrations were determined using a spectrophotometric assay with a 4:1 Free Glycerol Agent / Triglyceride Agent solution (Cat#: F6428 & T2449, Sigma, Free-Glycerol reagents and Triglyceride, St. Louis, MO) as previously described^45^.

### Real Time qPCR

We extracted total RNA from liver using Trizol (Life Technologies, Carlsbad, CA), which was reverse transcribed to cDNA with the M-MLV Reverse Transcriptase kit (Life Technologies). We performed RT-qPCR in a 7500 Fast Real Time PCR System (Applied Biosystems, Foster City, CA) using TaqMan Fast Universal PCR Master Mix (Life Technologies) and predesigned probes and primers for G6Pase and PEPCK from Applied Biosystems. Target gene expression was expressed as 2^−ΔΔCT^ by the comparative CT method^46^ and normalized to the expression of TATA-binding protein (Integrated DNA Technologies).

### GC-MS analysis

For analysis of bulk protein synthesis rates, we fractionated tissues according to our previously published procedures ^47–51^ including mitochondria, cytosolic, and mixed fractions. Skeletal muscles and liver tissues (20-30 mg) were homogenized 1:20 in isolation buffer (100 mM KCl, 40 mM Tris HCl, 10 mM Tris base, 5 mM MgCl2, 1 mM EDTA, and 1 mM ATP, pH = 7.5) with phosphatase and protease inhibitors (HALT, ThermoScientific, Rockford, IL, USA) using a bead homogenizer (Next Advance Inc., Averill Park, NY, USA). The subcellular fractions were then isolated via differential centrifugation and the protein pellets were isolated and purified, 250 μL of 1 M NaOH was added, and pellets were incubated for 15 min at 50°C. Proteins were hydrolyzed in 6 N HCl for 24 h at 120°C. To analyze the deuterium enrichment of alanine in the protein fractions we used Gas Chromatography-Mass Spectroscopy (7890A GC-Agilent 5975C MS, Agilent, Santa Clara, CA)^52,53^.

### Body water enrichment

To determine body water enrichment, 50 μl of plasma and 0–12% D_2_O standards (Sigma, 151882) were placed into the inner well of an o-ring screw cap and inverted on heating block overnight at 80°C. Samples were diluted 1:300 in ddH2O and analyzed on a liquid water isotope analyzer (Los Gatos Research, Los Gatos, CA, USA) against a standard curve prepared with samples having different concentrations of D2O (0-12%). The deuterium enrichments of both the protein (product) and the plasma (precursor) were used to calculate fraction new: Fraction new=E_product_/E_precursor_, where the E_product_ is the enrichment (E) of protein-bound alanine and E_precursor_ is the calculated maximum alanine enrichment from equilibration of the body water pool with alanine by MIDA.

### LC-MS/MS analysis

We used a combined targeted quantitative and kinetic proteomics as we previously described^54^. This approach analyzed two to five selected and validated peptides per mitochondrial protein in a series of panels that analyze proteins involved in the mitochondrial complexes as well as membrane-specific (inner and outer). The samples were analyzed using high resolution accurate mass (HRAM) measurements on a ThermoScientific Q-Exactive Plus hybrid quadrupole-orbitrap mass spectrometer at a m/z resolution of 140,000 configured with a splitless capillary column HPLC system (ThermoScientific Ultimate 3000). For protein quantification, the data were processed using Skyline^55^. The response for each protein was taken as the geometric mean of the response for the two to five peptides monitored for each protein. Changes in the abundance of the proteins were determined by normalization to the BSA internal standard. For analysis for kinetic proteomics, we used the D_2_Ome software package^56^, which analyzes the deuterium dependent change in the isotope pattern of each peptide in the full scan MS1 data. A database of mass, retention time, and sequence for each peptide of interest was assembled from LC-MS/MS peptide sequencing experiments on the same tissues. Mass spectral accuracy was confirmed by a <10% deviation from theory using peptides from the BSA internal standard^57^.

### Electroporation of DNA Construct

To perform imaging of mitochondrial structure, we electroporated the TA seven days prior to euthanasia with constructs to visualize the outer mitochondrial membrane and matrix. First, legs were shaved using clippers and Nair hair removal. While the rats were under isoflurane, we used an insulin syringe to inject 200 µl of 0.45 U/ml hyaluronidase (Sigma-Aldrich, St. Louis, MO, USA, #:H4272) into the TA muscle belly via three injections (from the distal, lateral, and proximal ends). Two hours after hyaluronidase injection, the rats were again briefly placed on isoflurane and were injected with a total of 40 µg of DNA, 5 µg pCAG-mtYFP^58^ and 5 µg of pCAG-TOM20tdTomato via three injections as described above. To electroporate the skeletal muscle, a small amount of Ultrasound Transmission Gel (Aquasonic 100) was placed on the TA. The Tweezertrodes were placed around the TA muscle where one side was located on top of the belly and the other was on the medial head of the gastrocnemius (gastroc). An electrical stimulus was applied with a ECM830 electroporator (model# 45-0662, BTX, Holliston, MA, USA) at a voltage of 86 V, ten pulses, a pulse length of 20 milliseconds, a distance between electrodes of 2 mm, and 0.5 milliseconds between pulses^58^.

### Muscle fixation and sectioning

Immediately following their excision, electroporated TA muscles were placed in 4% paraformaldehyde at 4°C overnight. The following morning, muscles were moved through a 4°C sucrose gradient protocol, starting with 2 hours in 10% sucrose in PBS, 2 hours in 20% sucrose in PBS, and lastly stored in 30% sucrose in PBS until they were further processed. Muscles were taken out of the 30% sucrose in PBS, pinned onto a piece of cork, and immediately frozen in freezing-cold isopentane. A ½ cm block was cut out of the frozen muscles and placed on a different piece of cork and embedded in OCT before being rapidly placed in freezing-cold isopentane for a minimum of 30 seconds. Ten micron thick muscle cross-sections were made using a Cryostar Cryotome and were mounted on slides. Slides were covered with ProLong™ Gold Antifade Mountant (Invitrogen P36930), a coverslip, and left to dry in a dark box at room temp for 24 hours. Slides were stored at −20°C until imaged.

### Imaging

We imaged a minimum of five fibers per muscle using a Zeiss LSM 880 with Airyscan FAST (Zeiss, Germany). Randomly selected electroporated fibers were imaged with a 40X oil immersion lens. Following the acquisition of images, the Zeiss ZEN Black program automatically processed the images. Composite images were semi-automatically analyzed using Fiji. We measured fiber cross-sectional area, the number of mitochondrial branches, and mitochondrial cross-sectional area. Additionally, we assessed the co-localization of the two fluorophores used to stain the mitochondrial outer membrane and the matrix as a proxy of cristae structure. Lastly, we further analyzed mitochondrial morphology using the same techniques by assessing the two unique mitochondrial populations in skeletal muscle, the intermyofibrillar and the subsarcolemmal, which we defined as the mitochondria located in the outer 10% of the fiber’s cross-section. Individual fibers were identified and measured using the polygon tool for fiber cross sectional area. Each fiber was automatically split by channel (red/green), segmented into four sections, and the brightness/contrast was automatically adjusted for uniformity and then fused back together. The threshold tool was automatically applied to both image channels and particle analysis was performed to assess mitochondrial number and cross-sectional area. The scaling interface identified the outer 10% of the fiber, and the same process was repeated to assess mitochondrial number, and mitochondrial cross-sectional area of the subsarcolemmal mitochondria^59^.

### High-resolution respirometry

To measure mitochondrial respiration and reactive oxygen species (ROS) production, we used our previously established high-resolution respirometry protocols (Oxygraph-2k, Oroboros Instruments, Innsbruck, Austria) on fiber bundles isolated from the middle third of the lateral head of the gastroc muscle^60–62^. For our protocol, Glutamate (10 mM), Malate (2 mM), and Pyruvate (5 mM) were added to determine the respiration and ROS production of CI. We then titrated ADP levels (0.125-11.8125 mM) to determine submaximal and maximal CI respiration. We added Cytochrome C (10 μm) to determine membrane integrity, followed by Succinate (10 mM) to determine CI+CII respiration (i.e. maximal oxidative phosphorylation) and ROS production. Maximal respiration was determined by uncoupling electron transfer capacity via the addition of the CCCP (1.5 μm) H+ ionophore. The addition of Rotenone (0.5 μm), a CI inhibitor, determined the respiration and ROS production of CII. To determine the acute effects of metformin on mitochondrial respiration, we spiked an Oxygraph-2k chamber with 7.5 μm of metformin, waited 15 minutes, and then repeated our mitochondrial respiration protocol on another fiber bundle from the MET-WO group.

### Statistical Analyses

We performed a two-way ANOVA (strain x treatment) using Graphpad Prism 10 (San Diego, CA, USA) to determine differences between groups and a Tukey post hoc analysis to determine where significant differences occurred unless otherwise stated. For water consumption we used a two-way repeated measures ANOVA (strain x time). For high resolution respirometry, we performed a one-way ANOVA for each substrate addition and a Tukey post hoc analysis to determine where significant differences occurred. Outliers as determined by Grubbs’ test (Alpha = 0.05) and removed from analyses. We presented the results as mean ± standard error (SE) unless otherwise stated with *p* values less than 0.05 considered to be significant unless otherwise specified. To visual spatial representation of our quantitative and kinetic proteomics, we used dot plots and ChimeraX^63^ software to generate figures using R programming language (3.4.0) and “colorbewer” and “ggplot2” packages^64,65^.

## Data availability

All the data generated or analyzed during this study are included in the published article and its Supplementary Information and Source Data files and are available from the corresponding author upon reasonable request.

Correspondence and requests for materials should be addressed to B.F.M.

## Code Availability

The software used in this study is publicly available and accessed without restriction. R package colorbrewer is available at https://cran.r-project.org/web/packages/RColorBrewer/index.html. R package ggplot2 v.3.3.6 is available at https://cran.r-project.org/web/packages/ggplot2/index.html. R Studio v.4.1.2 was used to format data for ChimeraX input and analyses. The code used in this study can be accessed at 10.6084/m9.figshare.25301230.

## Acknowledgements

Some images were created with BioRender.com and ChimeraX. We would like to acknowledge the Center for Biomedical Data Sciences at the Oklahoma Medical Research Foundation for the assistance with bioinformatics and figure generation. We thank Samantha J. McKee at The University of Toledo for expert phenotyping, care, and maintenance of the LCR/HCR rat colony.

## Ethics declarations

The authors declare no competing interests.

