## Extended Data Fig. 1 for "Metformin treatment results in distinctive skeletal muscle mitochondrial remodeling in rats with different intrinsic aerobic capacities"

### Slide 1
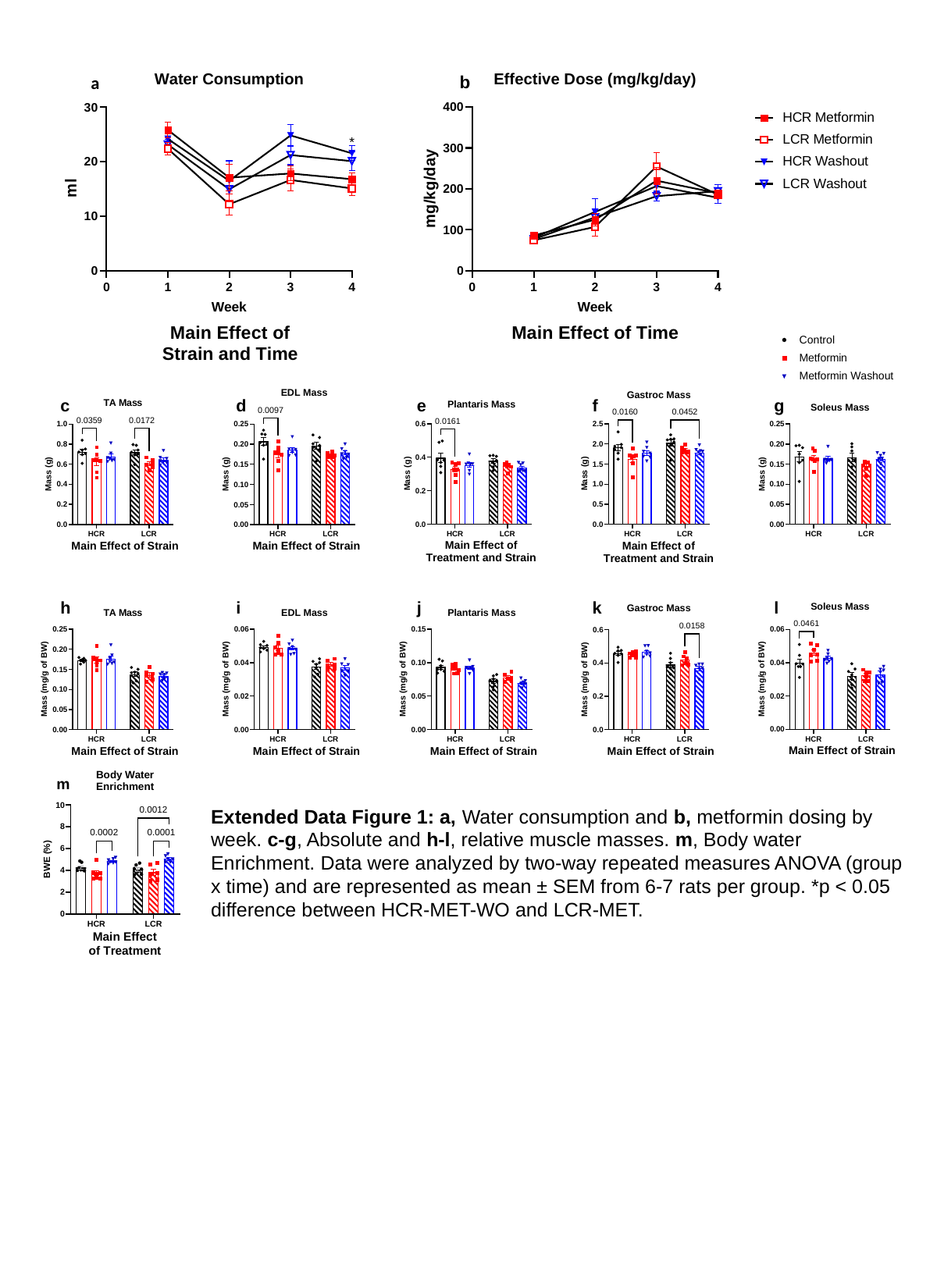

b
a
Extended Data Figure 1: a, Water consumption and b, metformin dosing by week. c-g, Absolute and h-l, relative muscle masses. m, Body water Enrichment. Data were analyzed by two-way repeated measures ANOVA (group x time) and are represented as mean ± SEM from 6-7 rats per group. *p < 0.05 difference between HCR-MET-WO and LCR-MET.
