## Extended Data Fig. 2 for "Metformin treatment results in distinctive skeletal muscle mitochondrial remodeling in rats with different intrinsic aerobic capacities"

### Slide 1
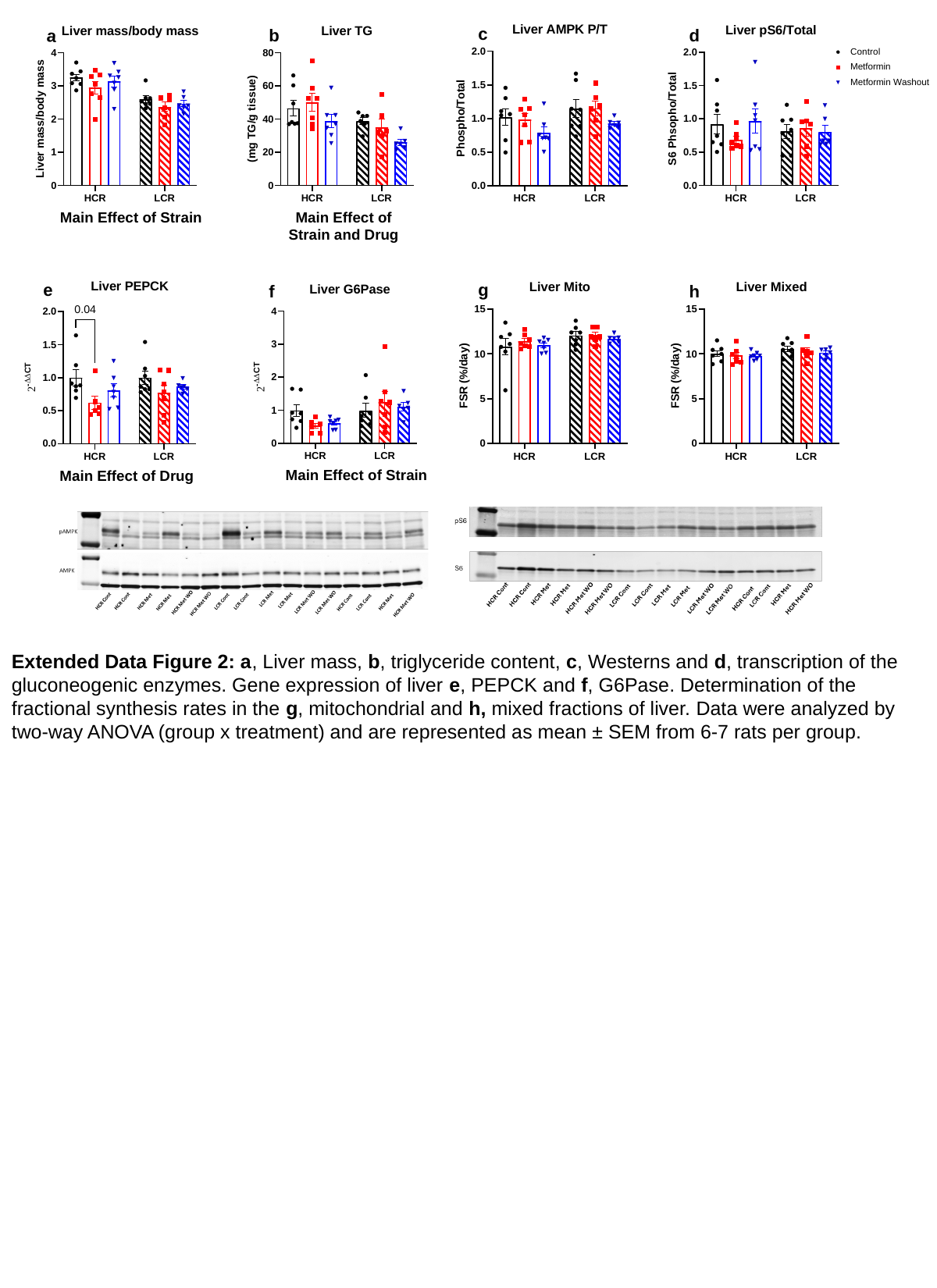

Extended Data Figure 2: a, Liver mass, b, triglyceride content, c, Westerns and d, transcription of the gluconeogenic enzymes. Gene expression of liver e, PEPCK and f, G6Pase. Determination of the fractional synthesis rates in the g, mitochondrial and h, mixed fractions of liver. Data were analyzed by two-way ANOVA (group x treatment) and are represented as mean ± SEM from 6-7 rats per group.
