## Extended Data Fig. 4 for "Metformin treatment results in distinctive skeletal muscle mitochondrial remodeling in rats with different intrinsic aerobic capacities"

### Slide 1
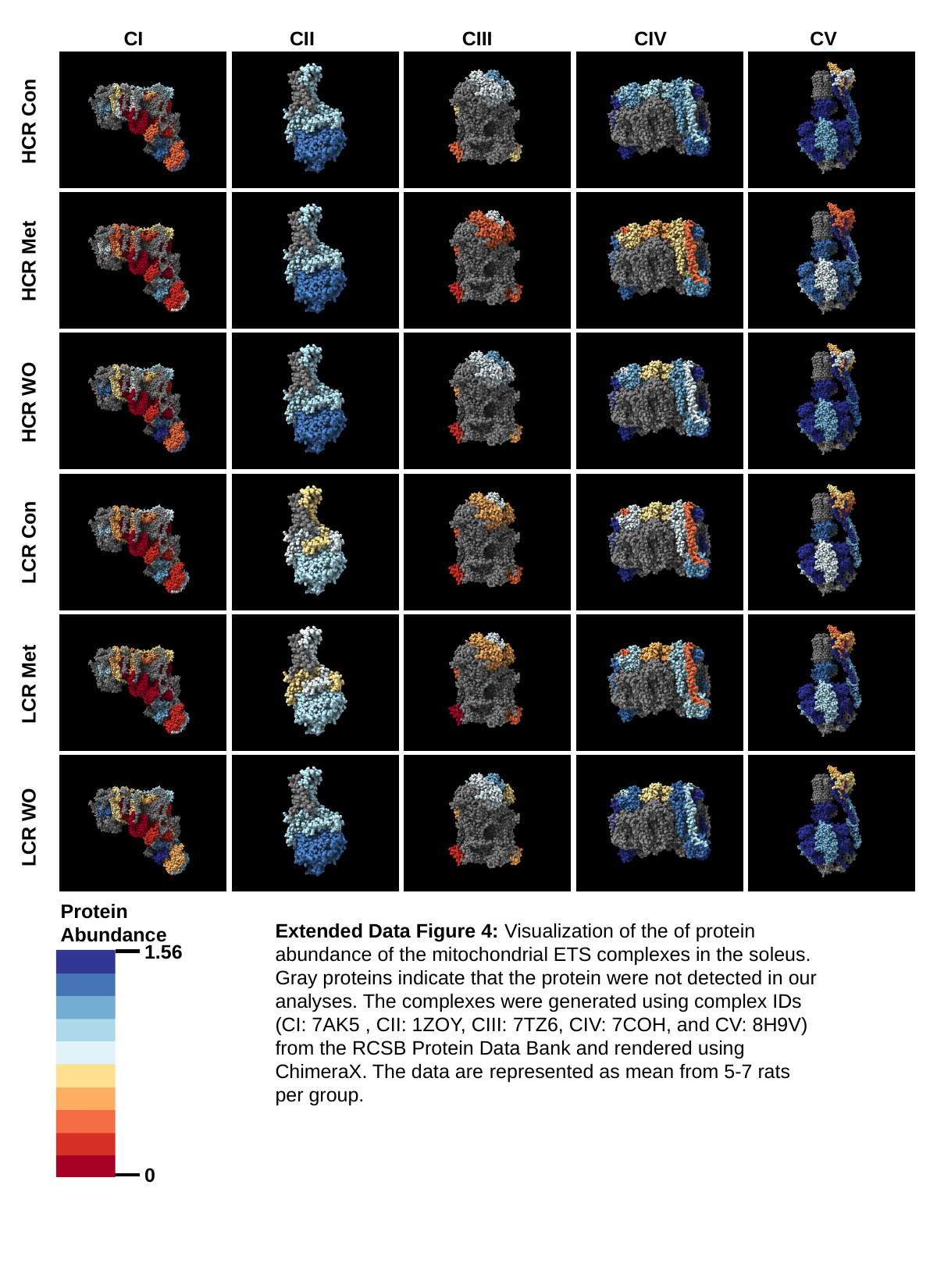

CI
CII
CIII
CIV
CV
HCR Con
HCR Met
HCR WO
LCR Con
LCR Met
LCR WO
Protein Abundance
1.56
0
Extended Data Figure 4: Visualization of the of protein abundance of the mitochondrial ETS complexes in the soleus. Gray proteins indicate that the protein were not detected in our analyses. The complexes were generated using complex IDs (CI: 7AK5 , CII: 1ZOY, CIII: 7TZ6, CIV: 7COH, and CV: 8H9V) from the RCSB Protein Data Bank and rendered using ChimeraX. The data are represented as mean from 5-7 rats per group.
