## Extended Data Fig. 5 for "Metformin treatment results in distinctive skeletal muscle mitochondrial remodeling in rats with different intrinsic aerobic capacities"

### Slide 1
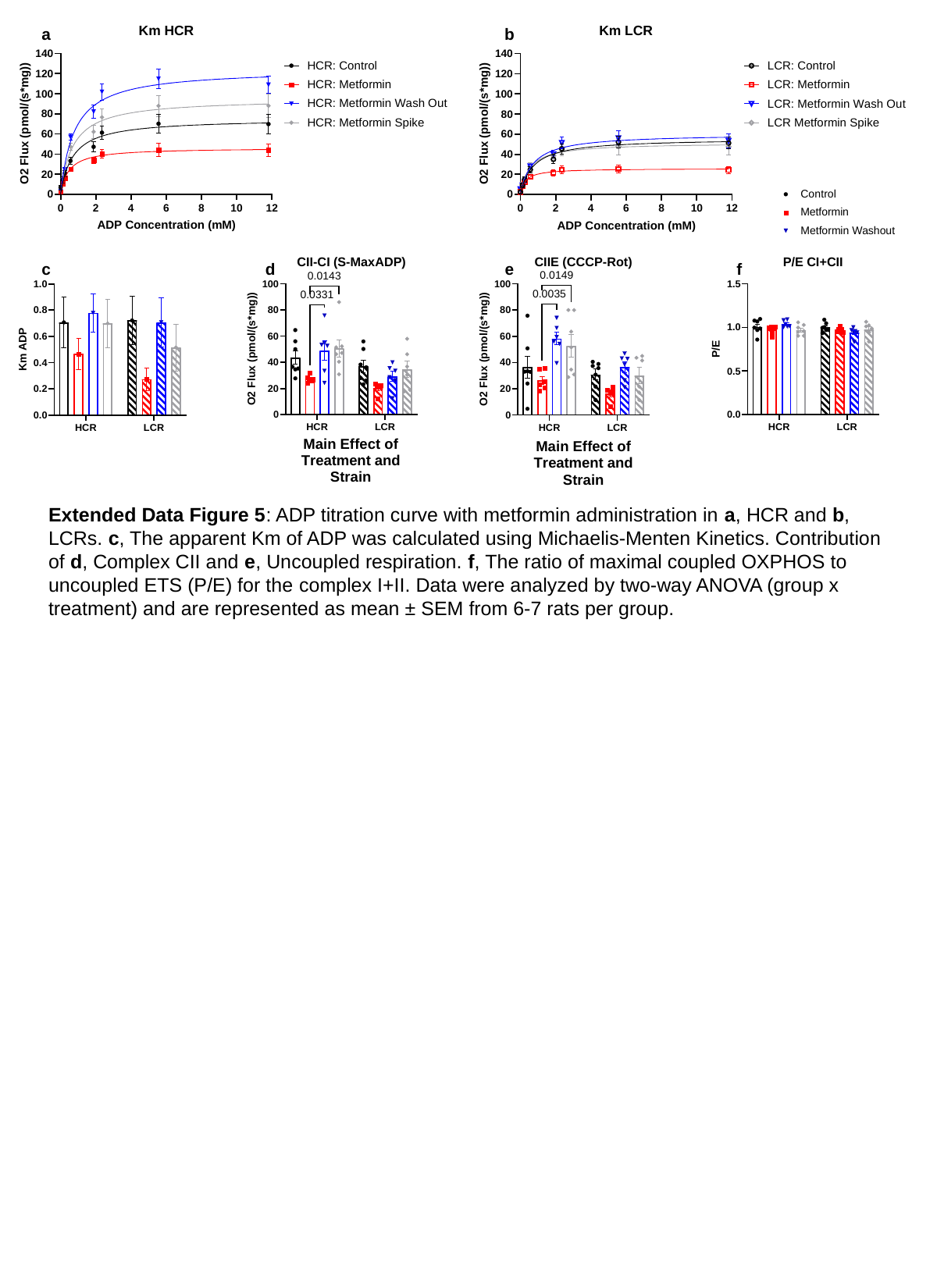

Extended Data Figure 5: ADP titration curve with metformin administration in a, HCR and b, LCRs. c, The apparent Km of ADP was calculated using Michaelis-Menten Kinetics. Contribution of d, Complex CII and e, Uncoupled respiration. f, The ratio of maximal coupled OXPHOS to uncoupled ETS (P/E) for the complex I+II. Data were analyzed by two-way ANOVA (group x treatment) and are represented as mean ± SEM from 6-7 rats per group.
