## Supplemental Table 1 for "Metformin treatment results in distinctive skeletal muscle mitochondrial remodeling in rats with different intrinsic aerobic capacities"

### Slide 1
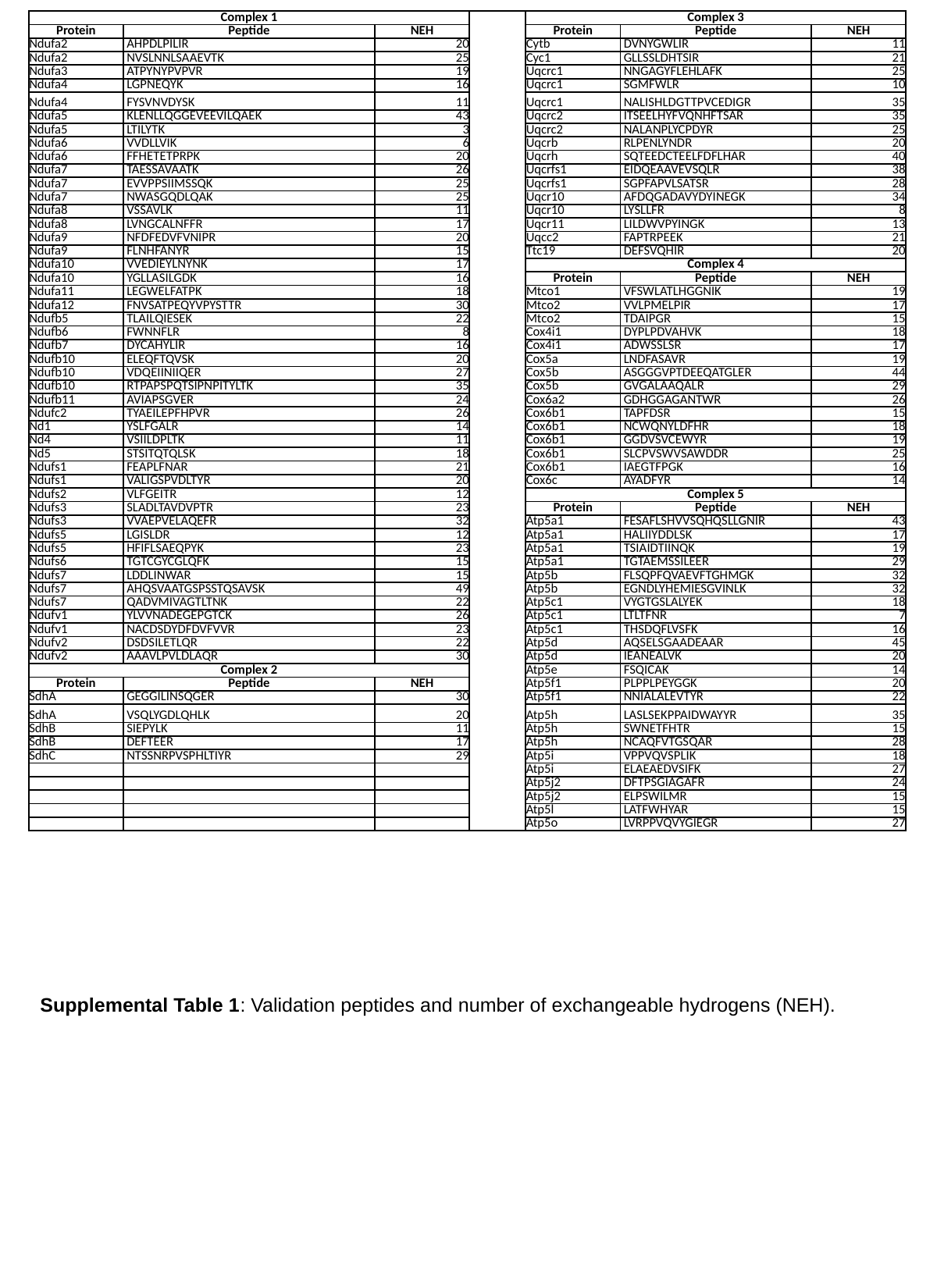

| Complex 1 | | | | Complex 3 | | |
| --- | --- | --- | --- | --- | --- | --- |
| Protein | Peptide | NEH | | Protein | Peptide | NEH |
| Ndufa2 | AHPDLPILIR | 20 | | Cytb | DVNYGWLIR | 11 |
| Ndufa2 | NVSLNNLSAAEVTK | 25 | | Cyc1 | GLLSSLDHTSIR | 21 |
| Ndufa3 | ATPYNYPVPVR | 19 | | Uqcrc1 | NNGAGYFLEHLAFK | 25 |
| Ndufa4 | LGPNEQYK | 16 | | Uqcrc1 | SGMFWLR | 10 |
| Ndufa4 | FYSVNVDYSK | 11 | | Uqcrc1 | NALISHLDGTTPVCEDIGR | 35 |
| Ndufa5 | KLENLLQGGEVEEVILQAEK | 43 | | Uqcrc2 | ITSEELHYFVQNHFTSAR | 35 |
| Ndufa5 | LTILYTK | 3 | | Uqcrc2 | NALANPLYCPDYR | 25 |
| Ndufa6 | VVDLLVIK | 6 | | Uqcrb | RLPENLYNDR | 20 |
| Ndufa6 | FFHETETPRPK | 20 | | Uqcrh | SQTEEDCTEELFDFLHAR | 40 |
| Ndufa7 | TAESSAVAATK | 26 | | Uqcrfs1 | EIDQEAAVEVSQLR | 38 |
| Ndufa7 | EVVPPSIIMSSQK | 25 | | Uqcrfs1 | SGPFAPVLSATSR | 28 |
| Ndufa7 | NWASGQDLQAK | 25 | | Uqcr10 | AFDQGADAVYDYINEGK | 34 |
| Ndufa8 | VSSAVLK | 11 | | Uqcr10 | LYSLLFR | 8 |
| Ndufa8 | LVNGCALNFFR | 17 | | Uqcr11 | LILDWVPYINGK | 13 |
| Ndufa9 | NFDFEDVFVNIPR | 20 | | Uqcc2 | FAPTRPEEK | 21 |
| Ndufa9 | FLNHFANYR | 15 | | Ttc19 | DEFSVQHIR | 20 |
| Ndufa10 | VVEDIEYLNYNK | 17 | | Complex 4 | | |
| Ndufa10 | YGLLASILGDK | 16 | | Protein | Peptide | NEH |
| Ndufa11 | LEGWELFATPK | 18 | | Mtco1 | VFSWLATLHGGNIK | 19 |
| Ndufa12 | FNVSATPEQYVPYSTTR | 30 | | Mtco2 | VVLPMELPIR | 17 |
| Ndufb5 | TLAILQIESEK | 22 | | Mtco2 | TDAIPGR | 15 |
| Ndufb6 | FWNNFLR | 8 | | Cox4i1 | DYPLPDVAHVK | 18 |
| Ndufb7 | DYCAHYLIR | 16 | | Cox4i1 | ADWSSLSR | 17 |
| Ndufb10 | ELEQFTQVSK | 20 | | Cox5a | LNDFASAVR | 19 |
| Ndufb10 | VDQEIINIIQER | 27 | | Cox5b | ASGGGVPTDEEQATGLER | 44 |
| Ndufb10 | RTPAPSPQTSIPNPITYLTK | 35 | | Cox5b | GVGALAAQALR | 29 |
| Ndufb11 | AVIAPSGVER | 24 | | Cox6a2 | GDHGGAGANTWR | 26 |
| Ndufc2 | TYAEILEPFHPVR | 26 | | Cox6b1 | TAPFDSR | 15 |
| Nd1 | YSLFGALR | 14 | | Cox6b1 | NCWQNYLDFHR | 18 |
| Nd4 | VSIILDPLTK | 11 | | Cox6b1 | GGDVSVCEWYR | 19 |
| Nd5 | STSITQTQLSK | 18 | | Cox6b1 | SLCPVSWVSAWDDR | 25 |
| Ndufs1 | FEAPLFNAR | 21 | | Cox6b1 | IAEGTFPGK | 16 |
| Ndufs1 | VALIGSPVDLTYR | 20 | | Cox6c | AYADFYR | 14 |
| Ndufs2 | VLFGEITR | 12 | | Complex 5 | | |
| Ndufs3 | SLADLTAVDVPTR | 23 | | Protein | Peptide | NEH |
| Ndufs3 | VVAEPVELAQEFR | 32 | | Atp5a1 | FESAFLSHVVSQHQSLLGNIR | 43 |
| Ndufs5 | LGISLDR | 12 | | Atp5a1 | HALIIYDDLSK | 17 |
| Ndufs5 | HFIFLSAEQPYK | 23 | | Atp5a1 | TSIAIDTIINQK | 19 |
| Ndufs6 | TGTCGYCGLQFK | 15 | | Atp5a1 | TGTAEMSSILEER | 29 |
| Ndufs7 | LDDLINWAR | 15 | | Atp5b | FLSQPFQVAEVFTGHMGK | 32 |
| Ndufs7 | AHQSVAATGSPSSTQSAVSK | 49 | | Atp5b | EGNDLYHEMIESGVINLK | 32 |
| Ndufs7 | QADVMIVAGTLTNK | 22 | | Atp5c1 | VYGTGSLALYEK | 18 |
| Ndufv1 | YLVVNADEGEPGTCK | 26 | | Atp5c1 | LTLTFNR | 7 |
| Ndufv1 | NACDSDYDFDVFVVR | 23 | | Atp5c1 | THSDQFLVSFK | 16 |
| Ndufv2 | DSDSILETLQR | 22 | | Atp5d | AQSELSGAADEAAR | 45 |
| Ndufv2 | AAAVLPVLDLAQR | 30 | | Atp5d | IEANEALVK | 20 |
| Complex 2 | | | | Atp5e | FSQICAK | 14 |
| Protein | Peptide | NEH | | Atp5f1 | PLPPLPEYGGK | 20 |
| SdhA | GEGGILINSQGER | 30 | | Atp5f1 | NNIALALEVTYR | 22 |
| SdhA | VSQLYGDLQHLK | 20 | | Atp5h | LASLSEKPPAIDWAYYR | 35 |
| SdhB | SIEPYLK | 11 | | Atp5h | SWNETFHTR | 15 |
| SdhB | DEFTEER | 17 | | Atp5h | NCAQFVTGSQAR | 28 |
| SdhC | NTSSNRPVSPHLTIYR | 29 | | Atp5i | VPPVQVSPLIK | 18 |
| | | | | Atp5i | ELAEAEDVSIFK | 27 |
| | | | | Atp5j2 | DFTPSGIAGAFR | 24 |
| | | | | Atp5j2 | ELPSWILMR | 15 |
| | | | | Atp5l | LATFWHYAR | 15 |
| | | | | Atp5o | LVRPPVQVYGIEGR | 27 |
Supplemental Table 1: Validation peptides and number of exchangeable hydrogens (NEH).
